# Sialylation of EGFR by ST6GAL1 induces receptor activation and modulates trafficking dynamics

**DOI:** 10.1101/2023.06.03.543566

**Authors:** Katherine E. Ankenbauer, Tejeshwar C. Rao, Alexa L. Mattheyses, Susan L. Bellis

## Abstract

Aberrant glycosylation is a hallmark of a cancer cell. One prevalent alteration is an enrichment in α2,6-linked sialylation of N-glycosylated proteins, a modification directed by the ST6GAL1 sialyltransferase. ST6GAL1 is upregulated in many malignancies including ovarian cancer. Prior studies have shown that the addition of α2,6 sialic acid to the Epidermal Growth Factor Receptor (EGFR) activates this receptor, although the mechanism was largely unknown. To investigate the role of ST6GAL1 in EGFR activation, ST6GAL1 was overexpressed in the OV4 ovarian cancer line, which lacks endogenous ST6GAL1, or knocked down in the OVCAR-3 and OVCAR-5 ovarian cancer lines, which have robust ST6GAL1 expression. Cells with high expression of ST6GAL1 displayed increased activation of EGFR and its downstream signaling targets, AKT and NFκB. Using biochemical and microscopy approaches, including Total Internal Reflection Fluorescence (TIRF) microscopy, we determined that the α2,6 sialylation of EGFR promoted its dimerization and higher order oligomerization. Additionally, ST6GAL1 activity was found to modulate EGFR trafficking dynamics following EGF-induced receptor activation. Specifically, EGFR sialylation enhanced receptor recycling to the cell surface following activation while simultaneously inhibiting lysosomal degradation. 3D widefield deconvolution microscopy confirmed that in cells with high ST6GAL1 expression, EGFR exhibited greater co-localization with Rab11 recycling endosomes and reduced co-localization with LAMP1-positive lysosomes. Collectively, our findings highlight a novel mechanism by which α2,6 sialylation promotes EGFR signaling by facilitating receptor oligomerization and recycling.

## Introduction

The receptor tyrosine kinase Epidermal Growth Factor Receptor (EGFR) has been the subject of intensive research due to its key roles in normal and aberrant epithelial cell physiology (1). During development and under normal physiological conditions, EGFR promotes cell survival and proliferation, and also regulates cell differentiation (2). Alterations in EGFR signaling are prevalent in many epithelial malignancies. EGFR and its ligands are commonly overexpressed in tumors, and moreover, EGFR frequently acquires mutations that drive constitutive receptor activation. This, in turn, promotes cell proliferation, angiogenesis, metastasis and chemoresistance (3, 4). As an example of a cancer-associated EGFR alteration, EGFRvIII, a truncated form of EGFR, has a mutated ectodomain that mediates ligand-independent receptor activation (5). Many other cancer types harbor EGFR variants with mutations in the intracellular domain that foster protein stability (6). An understanding of EGFR activation and signaling is crucial for the therapeutic targeting of this receptor in cancer treatment.

EGFR and its downstream signaling pathways are complex, and regulation occurs at multiple molecular levels. Under basal conditions, EGFR predominantly exists as an auto-inhibited monomer at the plasma membrane. However, when stimulated with EGF, the auto-inhibitory tether releases, facilitating receptor homodimerization, subsequent autophosphorylation of the cytosolic tails, and activation of intracellular signaling cascades such as PI3K/AKT/mTOR, Ras/Raf/MEKK/ERK, and NFκB (7–9). Following activation, EGFR is internalized and then trafficked to various subcellular compartments depending upon the context (7). For instance, EGFR can be ubiquitinated and shuttled to the lysosome, where it is degraded, or recycled back to the cell surface to promote further signaling. Where EGFR localizes following activation and internalization depends upon factors such as the type and concentration of EGFR ligands within the microenvironment (10, 11). The balance between EGFR degradation and recycling is a key mechanism controlling how much signal the cell receives.

Another important factor in EGFR regulation is its glycosylation state. EGFR is a highly N-glycosylated protein, containing 11 canonical N-glycosylation consensus sequences and 4 non-canonical sequences (12, 13). Evidence suggests that all 11 canonical, and 1 noncanonical, sites are glycosylated (14). Previous studies have shown that the N-glycosylation of EGFR is pivotal for its structure and function. N-glycans influence EGFR conformation, ligand binding capabilities, and the orientation of the EGFR ectodomain relative to the plasma membrane (15, 16). Furthermore, N-glycosylation at a specific site (Asn-579) plays an essential role in maintaining the auto-inhibitory tether present on EGFR monomers (17). Thus, the glycosylation of EGFR exerts another layer of regulation in EGFR signaling.

EGFR is aberrantly glycosylated in cancer cells due to alterations in the expression and activity of various glycosyltransferases. One such glycosyltransferase is the ST6GAL1 sialyltransferase, which is upregulated in numerous malignancies including ovarian cancer (18–21). ST6GAL1 adds an α2,6-linked sialic acid to the terminus of N-glycans on select glycoproteins including EGFR (22–27). We and others have shown that the α2,6 sialylation of EGFR activates this receptor (22–25), however inhibitory effects of sialylation have also been reported (26–29). Furthermore, our group determined that the ST6GAL1-mediated sialylation of EGFR promotes epithelial-to-mesenchymal transition (22), resistance to the tyrosine kinase inhibitor, gefitinib (23), and mechanotransduction (24). These results point to a seminal role for EGFR sialylation in cancer cell behavior, however the molecular mechanisms by which α2,6 sialylation regulates EGFR dynamics and downstream signaling remain largely unknown. In the present study, we report that ST6GAL1-mediated sialylation activates EGFR in seven different cancer cell models including ovarian, pancreatic and colon cancer cells. To interrogate the mechanism of receptor activation, ST6GAL1 was overexpressed in the OV4 ovarian cancer line, which lacks endogenous ST6GAL1, or knocked-down in OVCAR-3 and OVCAR-5 ovarian cancer cells, which have high levels of ST6GAL1. Results from these models suggest that α2,6 sialylation of EGFR facilitates receptor dimerization and higher order clustering, leading to increased receptor activation and downstream signaling through AKT and NFκB. Additionally, the sialylation of EGFR by ST6GAL1 promotes recycling of the receptor to the cell surface, while preventing degradation. Taken together, these results highlight a novel glycosylation-dependent mechanism by which cancer cells hijack EGFR signaling to enhance tumor-promoting signaling pathways.

## Results

### Cells with high levels of ST6GAL1 exhibit greater EGF-dependent activation of EGFR

To investigate the effects of ST6GAL1-mediated sialylation on EGFR activity, we assessed EGFR activation in pancreatic, ovarian and colon cancer cell lines in which ST6GAL1 expression was directly modulated. The pancreatic cancer cell lines, MiaPaCa-2, S2-LM7AA, and S2-013, as well as the ovarian cancer cell lines, OVCAR-3 and OVCAR-5, have substantial ST6GAL1 expression, typical of most cancer cells. Accordingly, ST6GAL1 expression was knocked-down (KD) in these lines (Fig. 1A). As controls, cells were transduced with either a non-targeting shRNA sequence (shC) or an empty vector (EV) construct. Conversely, ST6GAL1 was overexpressed (OE) in the OV4 ovarian, and SW48 colon cancer lines, which have unusually low levels of endogenous ST6GAL1 (Fig. 1B). EV cells served as the control. The cell lines were then treated with 100 ng/mL EGF for 15 minutes and EGFR activation was monitored by immunoblotting for phosphorylated EGFR (p-EGFR, pY1068). All of the cell lines with ST6GAL1 KD had diminished EGF-induced EGFR activation relative to controls (Fig. 1C), whereas the OV4 and SW48 lines with ST6GAL1 OE had enhanced EGFR activation compared with EV cells (Fig. 1D). These data show that α2,6 sialylation consistently activates EGFR in a wide range of cancer cell models, despite differences in genetic backgrounds or organ site. Furthermore, α2,6 sialylation activates EGFR in the SW48 cell model, which reportedly has an EGFR mutation (G719S) that causes ligand-independent receptor activation (30).

**FIGURE 1.**
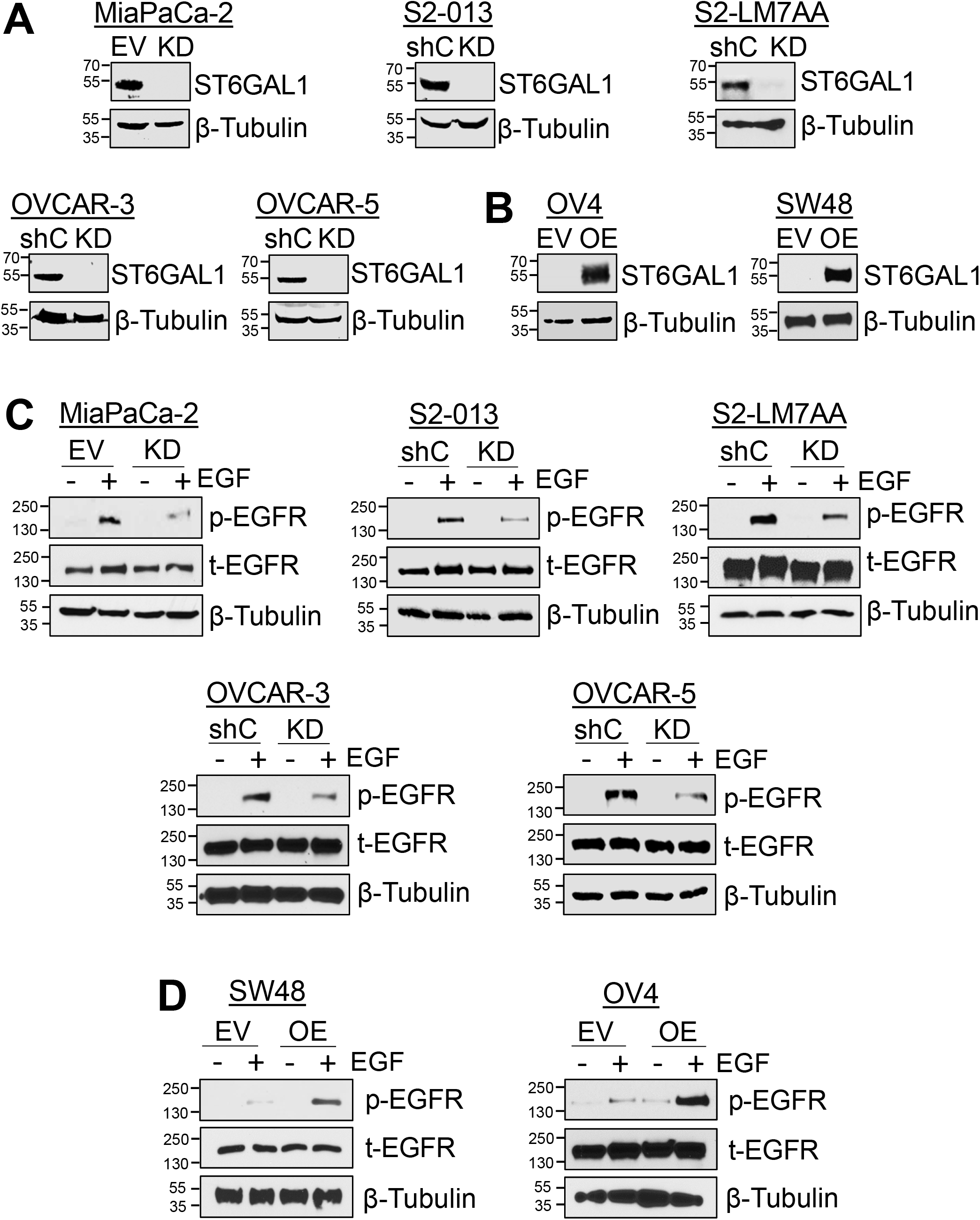
ST6GAL1-mediated sialylation of EGFR promotes its activation in multiple cell models. A). ST6GAL1 was stably knocked-down (KD) in cells with high endogenous ST6GAL1 expression (MiaPaCa-2, S2-013, S2-LM7AA, OVCAR-3, and OVCAR-5) using lentivirus encoding an shRNA sequence targeting ST6GAL1. As controls, cells were either transduced with lentivirus containing shRNA targeting GFP (shC) or with an empty vector (EV) construct. B). Cells with undetectable endogenous ST6GAL1 (SW48 and OV4) were stably transduced with ST6GAL1-encoding cDNA to overexpress (OE) the enzyme, or with an EV construct. All cell lines represent polyclonal populations. C). Cells with or without ST6GAL1 KD were treated with 100 ng/mL EGF for 15 minutes and immunoblotted for p-EGFR (pY1068) and total EGFR (t-EGFR). D). Cells with or without ST6GAL1 OE were treated with 100 ng/mL EGF for 15 minutes and immunoblotted for p-EGFR and t-EGFR.

### ST6GAL1-mediated sialylation does not alter the overall expression of EGFR or capacity of EGFR to bind ligand

To elucidate the molecular pathways by which ST6GAL1 regulates EGFR activation, we performed mechanistic studies using the three ovarian cancer cell lines, OV4, OVCAR-3 and OVCAR-5. We first confirmed that the modulation of ST6GAL1 expression led to a concomitant change in surface α2,6 sialylation. Cells were stained with *Sambucus nigra agglutinin* (SNA), a lectin that binds specifically to α2,6 sialic acids, and analyzed by flow cytometry. OV4 OE cells had increased surface levels of α2,6 sialic acid compared to EV cells, while OVCAR-3 and OVCAR-5 KD cells had reduced α2,6 sialylation compared to shC controls (Fig. 2A). We then verified that EGFR was a direct target for α2,6 sialylation, as has been previously reported (22–27). To this end, α2,6 sialylated proteins were precipitated using SNA-agarose and the precipitates were immunoblotted for EGFR. OV4 OE cells had higher levels of α2,6 sialylated EGFR, whereas OVCAR-3 and OVCAR-5 KD cells had decreased levels of α2,6 sialylated EGFR, relative to their respective controls (Fig. 2B). Immunoblots of whole cell lysates used as inputs for SNA precipitation showed that modulating ST6GAL1 expression did not alter EGFR protein expression (Fig. 2B). We also measured basal levels of EGFR on the cell surface by flow cytometry. Cells with differential expression of ST6GAL1 had comparable levels of surface EGFR (Fig. 2C). To determine if α2,6 sialylation of EGFR affected ligand binding, cells were incubated with EGF concentrations ranging from 0.39 nM to 200 nM and EGF binding was quantified by flow cytometry to create a ligand binding curve. No significant differences were detected in the capacity of sialylated EGFR to bind EGF (Fig. 2D).

**FIGURE 2.**
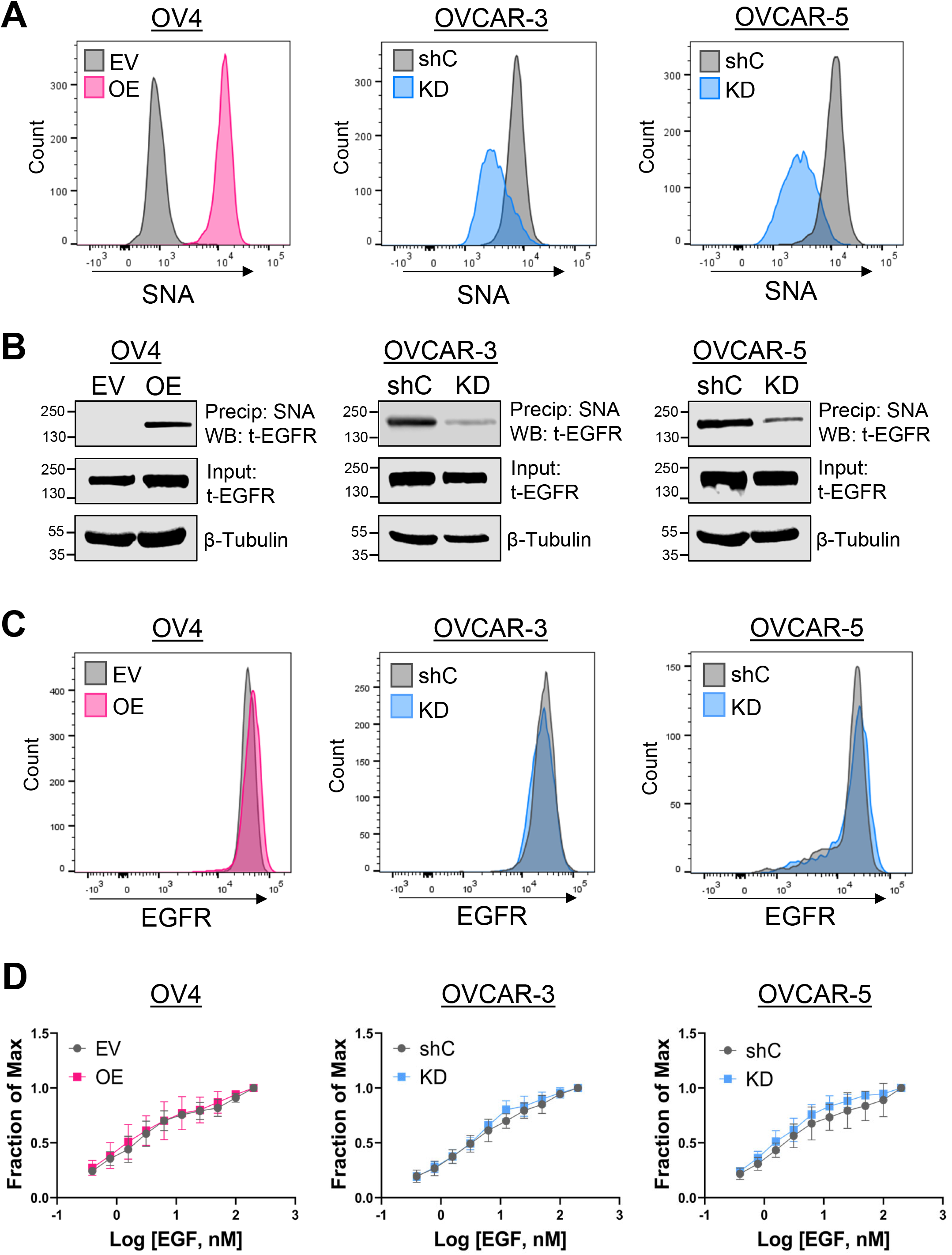
α2,6 sialylation of EGFR does not affect EGFR expression or ligand binding. A). Levels of α2,6 sialylation on the cell surface were assessed by staining cells with SNA and measuring via flow cytometry. B). Cell lysates were precipitated (Precip) with SNA-conjugated agarose and Western blotted (WB) for EGFR to determine the amount of α2,6 sialylated EGFR. Total EGFR expression was assessed by immunoblotting whole cell lysates (Input). C). Basal cell surface expression of EGFR was evaluated by flow cytometry. D). EGF binding was assessed using serial dilutions of EGF-biotin followed by flow cytometry. The x-axis depicts the log of the concentration of EGF and the y-axis is the fraction of maximal binding. Values from three independent experiments are graphed as the mean +/- S.D.

### Levels of α2,6 sialylation directly correlate with EGFR activation

To further corroborate the sialylation-dependent activation of EGFR, we evaluated EGFR phosphorylation in cells with high or low levels of surface α2,6 sialylation. Wild-type OVCAR-3 and OVCAR-5 cells were used for these experiments because they naturally possess a range of α2,6 sialylation levels. OV4 cells were not included because they lack detectable expression of endogenous ST6GAL1. We first optimized a flow cytometry protocol for intracellular staining of p-EGFR. OVCAR-3 and OVCAR-5 cells were treated with or without EGF for 10 minutes to activate EGFR, and then permeabilized cells were incubated with antibody against p-EGFR (pY1068). As expected, EGF treatment increased the levels of p-EGFR (Fig. 3A). Next, we co-stained cells with SNA and anti-p-EGFR. OVCAR-3 and OVCAR-5 cells were gated for the 10% of cells with the highest levels of surface α2,6 sialylation, and the 10% with the lowest levels of α2,6 sialylation, referred to as “SNA high” and “SNA low” respectively (schematic in Fig. 3B). The levels of p-EGFR in the SNA high and SNA low populations for OVCAR-3 (Fig. 3C-D) and OVCAR-5 (Fig. 3E-F) cells were quantified by obtaining the mean fluorescent intensity (MFI). Importantly, SNA high cells had significantly greater activation of EGFR as compared with SNA low cells both in the presence and absence of EGF treatment. These data indicate that high levels of ST6GAL1-mediated sialylation strongly correlate with an increase in EGFR activation.

**FIGURE 3.**
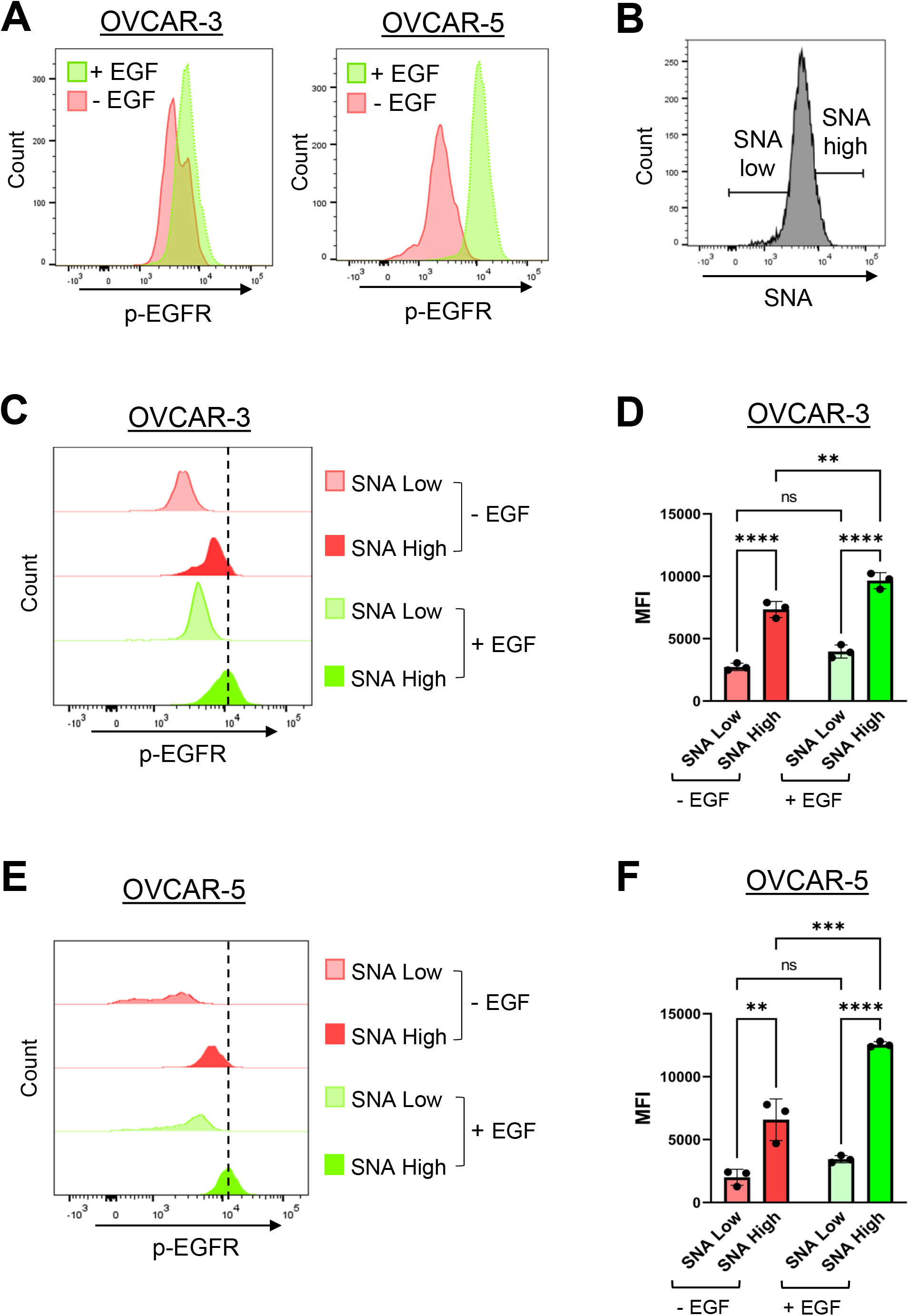
Cells with high levels of α2,6 sialylation have increased EGFR activation. OVCAR-3 and OVCAR-5 cells were treated with 100 ng/mL of EGF for 10 minutes, fixed, permeabilized, stained with SNA and/or p-EGFR, and then analyzed by flow cytometry. A). Histograms depicting intracellular staining for p-EGFR before and after treatment with EGF. B). Schematic of the gating strategy. The 10% of cells with the lowest levels of α2,6 sialylation were designated as “SNA low”, and the 10% of cells with the highest levels of α2,6 sialylation were designated as “SNA high”. SNA high and SNA low cells were assessed for levels of p-EGFR. C-D). p-EGFR levels in OVCAR-3 SNA high and SNA low cells. Representative experiment in (C) and quantification in (D). E-F). p-EGFR levels in OVCAR-5 SNA high and SNA low cells. Representative experiment in (E); quantification in (F). Dotted lines indicate the highest peak of the histograms. Graphs depict the MFI +/- S.D. from three independent experiments. (not significant, ns: p > 0.05, *: p < 0.05, **: p < 0.01, ***: p < 0.001, ****: p < 0.0001) as measured by a two-way ANOVA followed by Šidák’s multiple comparison test.

### Sialylation of EGFR enhances EGFR-mediated activation of AKT and NFκB p65, but not ***ERK***

The activation of EGFR stimulates multiple downstream signaling molecules including AKT, NFκB, and ERK. To determine the effects of sialylation on EGFR signaling, OV4 cells were treated with EGF for 5, 15, and 30 minutes and evaluated for p-EGFR (pY1068). OE cells had higher levels of activated EGFR than EV cells (representative blot in Fig. 4A, quantification in Fig. 4B). Correspondingly, OE cells exhibited enhanced activation of AKT (Fig. 4C-D) and NFκB p65 (Fig. 4E-F). Intriguingly, no differences were noted in ERK activation in EV vs. OE cells (Fig. 4G-H).

**FIGURE 4.**
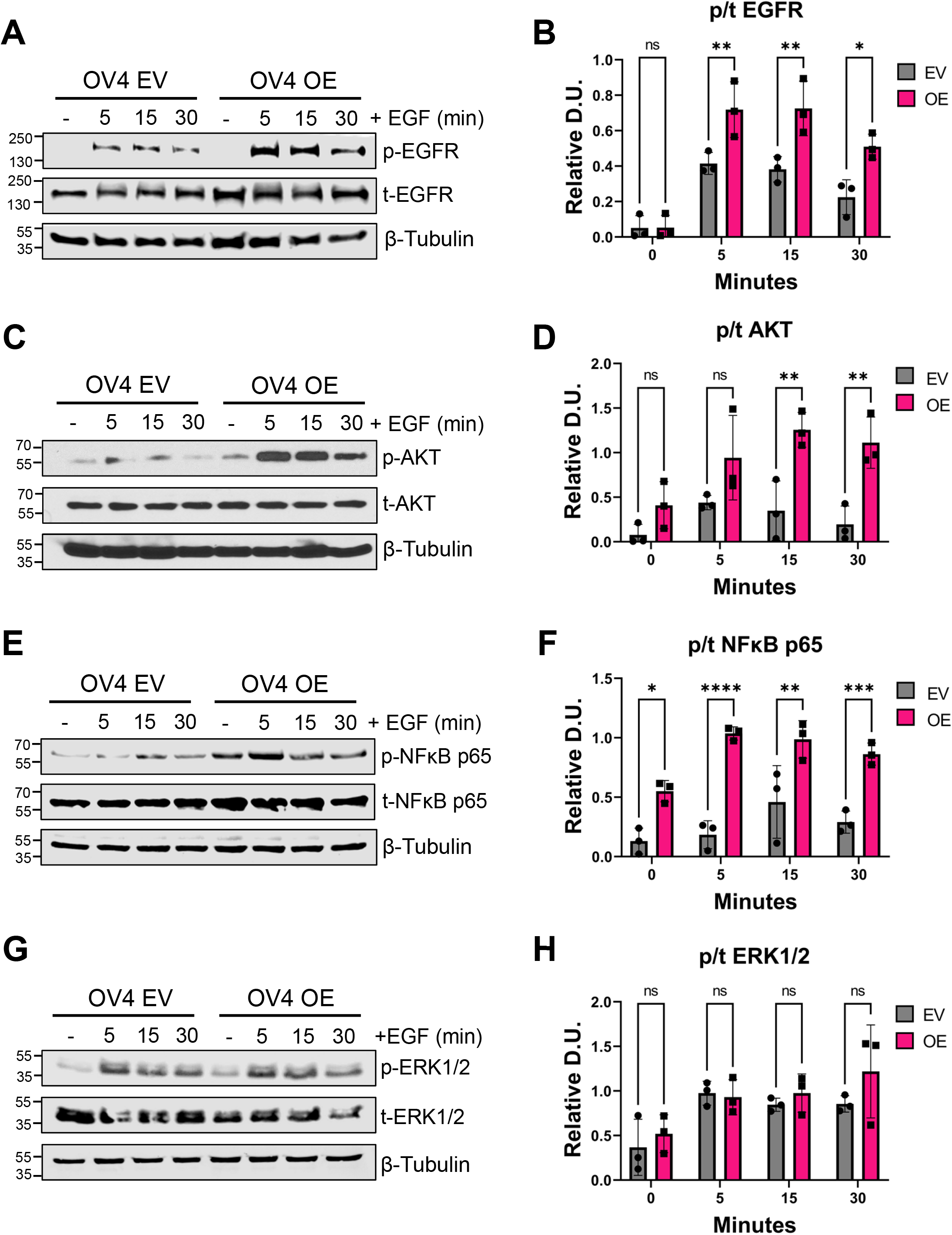
Overexpression of ST6GAL1 in OV4 cells activates EGFR, AKT and NFκB p65, but not ERK1/2. OV4 cells were treated with 100 ng/mL of EGF for 5, 15, or 30 minutes, or left untreated (-), and then cell lysates were immunoblotted for signaling molecules. A-B). Representative blot (A) and quantification (B) of p-EGFR and t-EGFR. C-D). Representative blot (C) and quantification (D) of p-AKT and t-AKT. E-F). Representative blot (E) and quantification (F) of p-NFκB p65 and t-NFκB p65. G-H). Representative blot (G) and quantification (H) of p-ERK1/2 and t-ERK1/2. Blots from three independent cell lysates were analyzed by densitometry and the phospho to total ratio (p/t) was calculated and normalized to β-tubulin. D.U. = densitometry units. Statistics were calculated using a two-way ANOVA followed by Šidák’s multiple comparison test. (ns: p > 0.05, *: p < 0.05, **: p < 0.01, ***: p < 0.001, ****: p < 0.0001).

Similar experiments were conducted with OVCAR-3 (Fig. 5) and OVCAR-5 (Fig. 6) cells with comparable results. In both cell models, ST6GAL1 KD decreased the activation of EGFR, AKT and NFκB p65, but did not alter signaling by ERK. Of note, in OVCAR-5 cells, EGF treatment had little effect on ERK activation, which may relate to the fact that OVCAR-5 cells have a KRAS G12V mutation (31).

**FIGURE 5.**
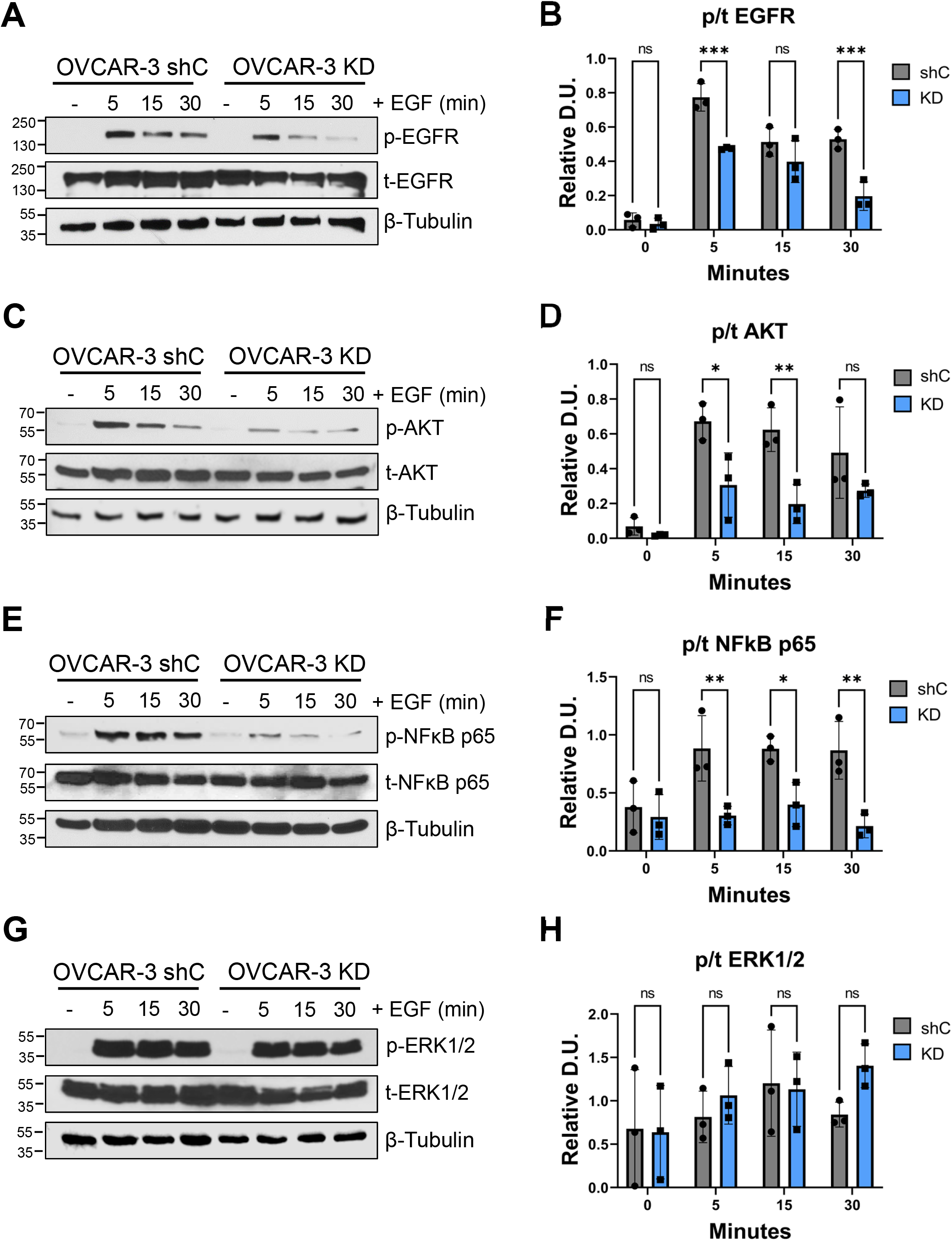
Knock-down of ST6GAL1 in OVCAR-3 cells diminishes activation of EGFR, AKT and NFκB p65, but does not affect ERK1/2 activation. OVCAR-3 cells were treated with 100 ng/mL of EGF for 5, 15, or 30 minutes or left untreated (-) and then cell lysates were immunoblotted for signaling molecules. A-B). p-EGFR and t-EGFR. C-D). p-AKT and t-AKT. E-F). p-NFκB p65 and t-NFκB p65. G-H). p-ERK1/2 and t-ERK1/2. The phospho to total ratio (p/t) was calculated and normalized to β-tubulin (“Relative D.U.”). Graphs depict the mean +/- S.D. for three independent immunoblots for each signaling molecule. Statistics were calculated using a two-way ANOVA followed by Šidák’s multiple comparison test. (ns: p > 0.05, *: p < 0.05, **: p < 0.01, ***: p < 0.001, ****: p < 0.0001).

**FIGURE 6.**
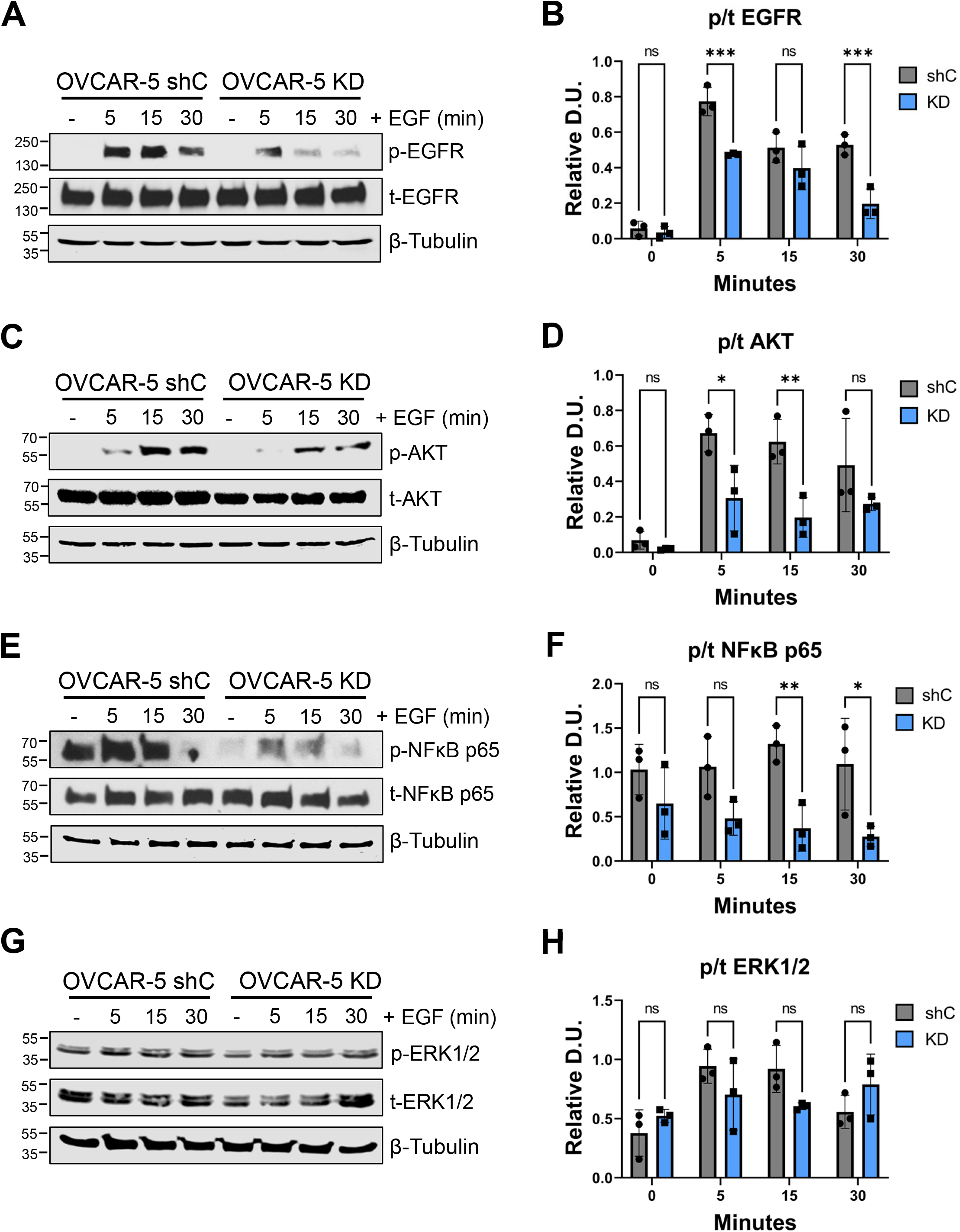
Knock-down of ST6GAL1 in OVCAR-5 cells diminishes activation of EGFR, AKT and NFκB p65, but does not affect ERK1/2 activation. OVCAR-5 cells were treated with 100 ng/mL of EGF for 5, 15, or 30 minutes or left untreated (-), and then cell lysates were immunoblotted for signaling molecules. A-B). p-EGFR and t-EGFR. C-D). p-AKT and t-AKT. E-F). p-NFκB p65 and t-NFκB p65. G-H). p-ERK1/2 and t-ERK1/2. The phospho to total ratio (p/t) was calculated and normalized to β-tubulin (“Relative D.U.”). Graphs depict the mean +/- S.D. for three independent immunoblots for each signaling molecule. Statistics were calculated using a two-way ANOVA followed by Šidák’s multiple comparison test. (ns: p > 0.05, *: p < 0.05, **: p < 0.01, ***: p < 0.001, ****: p < 0.0001).

### ST6GAL1-mediated sialylation of EGFR enhances EGFR homodimer formation

We next assessed the formation of the EGFR homodimer, a critical step in the activation of EGFR and downstream signaling pathways (7). To monitor homodimerization, we adapted a protocol from Turk et al. 2015 (32), in which surface homodimers are stabilized using the bis[sulfosuccinimidyl] suberate (BS^3^) cross-linking reagent. The presence of dimers on the cell surface was evaluated by immunoblotting, followed by densitometric quantification of the dimer to monomer ratio. In the OV4 cell line, significantly more dimer formation was observed in OE vs. EV cells in the absence of EGF, as well as following a 5-minute EGF treatment (Fig. 7A-B). In contrast, knockdown of ST6GAL1 in OVCAR-3 and OVCAR-5 cells led to a significant decrease in dimer formation, particularly at the early time points (Fig. 7C-F).

**FIGURE 7.**
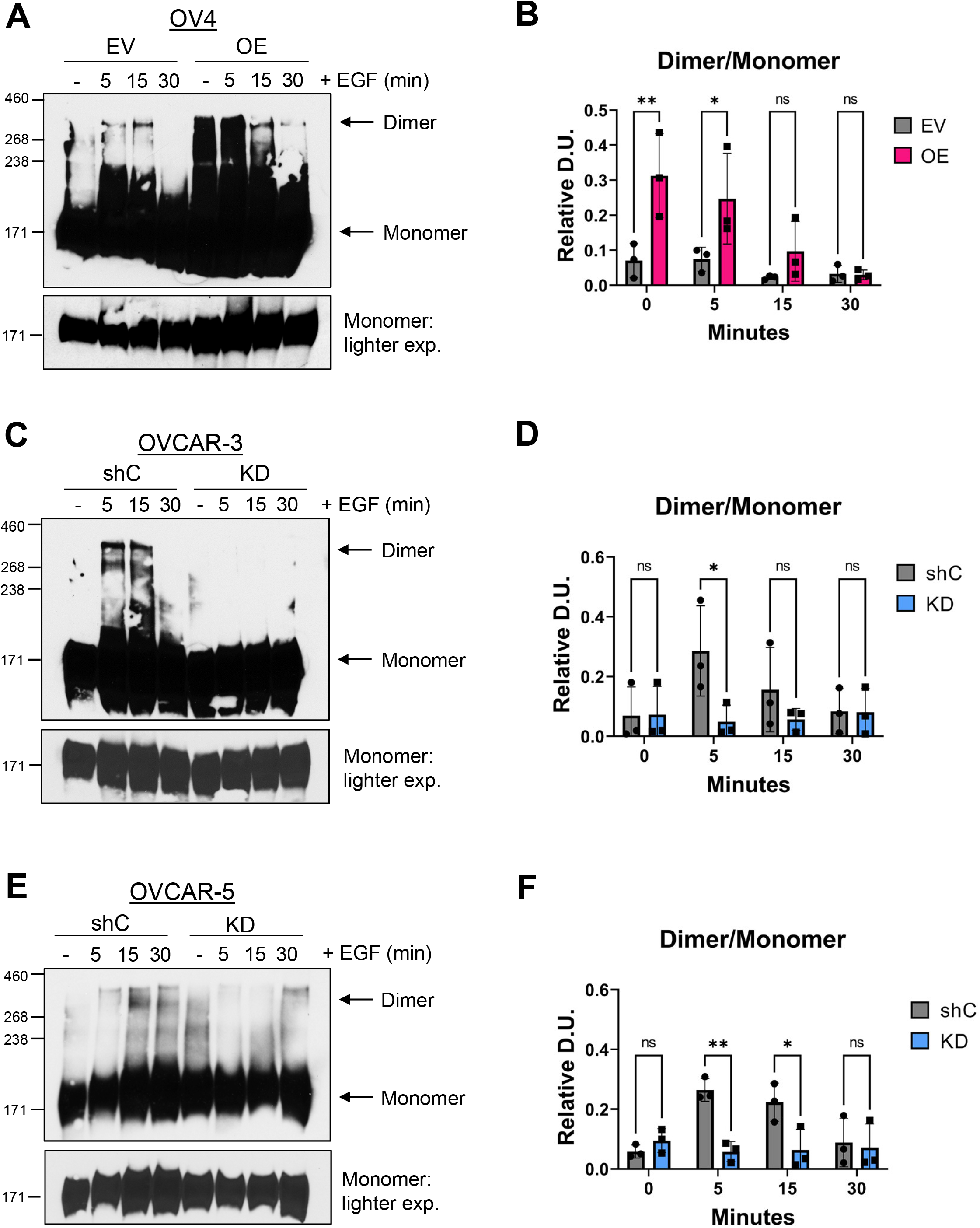
EGFR homodimer formation is enhanced by ST6GAL1-mediated sialylation. Cells were treated with 100 ng/mL of EGF for 5, 15, or 30 minutes, or left untreated (-). After treatment, proteins were crosslinked using 3 mM of BS^3^. Lysates were immunoblotted for EGFR. A high molecular weight ladder was used to distinguish monomers (∼170-180 kDa) from dimers (∼340-360 kDa). A-B). OV4 cells: representative immunoblot (A) and quantification (B) of the dimer to monomer ratio. A lighter exposure of the monomers (lower panel in A) was used for densitometric analyses. C-D). OVCAR-3 cells: representative immunoblot (C) and quantification (D) of the dimer to monomer ratio. E-F) OVCAR-5 cells: representative blot (E) and quantification (F) of the dimer to monomer ratio. Blots were analyzed by densitometry and the dimer to monomer ratio was calculated. Graphs depict mean +/- S.D. from three independent experiments. Statistics were calculated using a two-way ANOVA followed by Šidák’s multiple comparison test. (ns: p > 0.05, *: p < 0.05, **: p < 0.01).

### α2,6 sialylation of EGFR promotes receptor recycling to the cell surface

Following EGFR activation, EGFR internalizes into the early endosome, and then can either recycle back to the cell surface or translocate to the lysosome for degradation (7, 10). Accordingly, we evaluated the effects of ST6GAL1-mediated sialylation on EGFR recycling. Cells were first treated with cycloheximide (CHX) to prevent nascent EGFR synthesis (33). The levels of EGFR on the cell surface were then measured by flow cytometry for untreated cells, or for cells treated with EGF for 15 minutes to stimulate EGFR internalization. As expected, EGF treatment induced EGFR internalization, as indicated by the leftward peak shift (samples labeled as “0 min. recycling” in Fig. 8A, C, E). The amount of EGFR remaining on the cell surface following the 15-minute EGF treatment was designated as time 0. The EGF-containing media was then replaced with EGF-free media and cells were incubated for an additional 60 minutes to allow EGFR recycling to the cell surface (samples labeled as “60 min. recycling”). The percent recycling was calculated by comparing surface EGFR levels at the end of the 60-minute recycling period with the levels of surface EGFR at time 0. OV4 OE cells displayed significantly more EGFR recycling than EV cells (Fig. 8A,B), whereas ST6GAL1 KD in OVCAR-3 and OVCAR-5 cells diminished EGFR recycling (Fig. 8C-F). To confirm that the changes in EGFR levels at the cell surface were due to recycling, cells were treated with a recycling inhibitor, monensin (34). Treatment with monensin diminished EGFR recycling in cells with high ST6GAL1 while having a negligible effect on cells with low ST6GAL1 (Fig. S1). These data support the hypothesis that α2,6 sialylation of EGFR contributes to receptor recycling.

**FIGURE 8.**
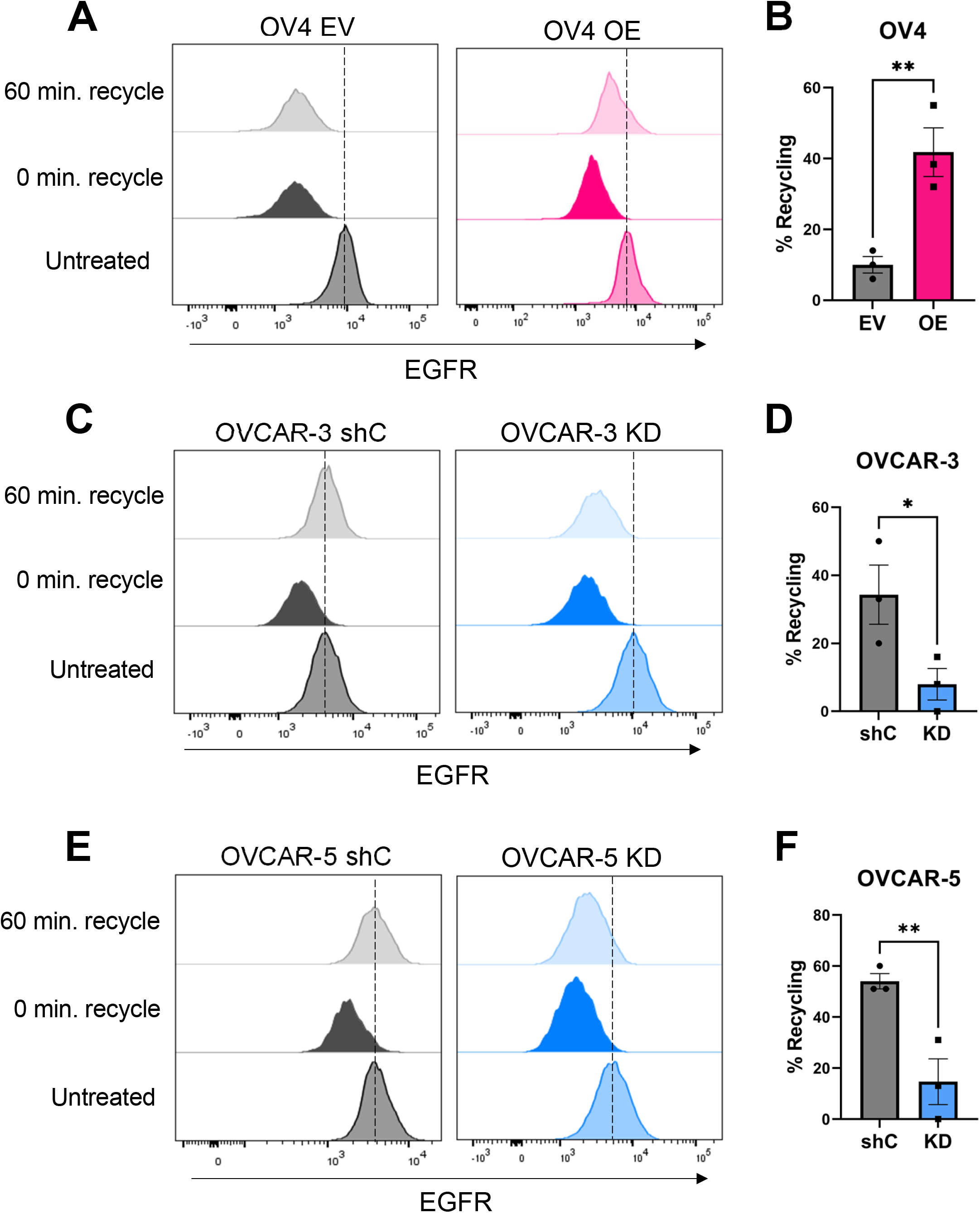
ST6GAL1-mediated sialylation of EGFR promotes receptor recycling to the cell surface. Cells were pretreated with 10 µg/mL of CHX to prevent nascent protein synthesis and then treated with 100 ng/mL of EGF for 15 minutes to induce EGFR internalization. At the end of this incubation, an aliquot of cells was fixed and analyzed for surface EGFR to obtain a baseline measurement immediately after the internalization step, designated as 0 minutes. The remaining cells were placed in EGF-free media and incubated for another 60 minutes at 37⁰C to allow EGFR recycling. These cells were then fixed and analyzed for surface EGFR. Cells untreated with EGF were used as a control. Percent recycling was calculated by comparing the MFI at 0 minutes to the MFI at 60 minutes. A-B). OV4 cells: representative histogram (A) and quantification (B) of EGFR recycling. C-D). OVCAR-3 cells: representative histogram (C) and quantification (D) of recycling. E-F). OVCAR-5 cells: representative histogram (E) and quantification (F) of EGFR recycling. Dotted lines indicate the peak MFI of untreated cells. Graphs depict mean and S.D. from three independent experiments. Statistics were calculated using a Student’s t-test (*: p < 0.05, **: p < 0.01).

### Sialylation of EGFR protects against EGFR degradation following EGF treatment

We next evaluated the effects of α2,6 sialylation on EGFR degradation following EGF treatment. Cells were pre-treated with CHX to prevent nascent EGFR synthesis and then incubated with EGF over a 120-minute interval. As controls, cells were either left untreated, or treated for 120 minutes with CHX alone (to assess the amount of EGFR degradation in the absence of EGF). Notably, OV4 cells with ST6GAL1 OE exhibited minimal EGF-stimulated EGFR degradation over the 120-minute incubation, while substantial degradation was observed in EV cells (Fig. 9A-B). No differences were noted in the levels of EGFR in the absence of EGF treatment or in the presence of CHX alone, confirming that EGFR degradation was secondary to the effects of EGF stimulation. Consistent with results from OV4 cells, OVCAR-3 and OVCAR-5 cells with ST6GAL1 KD exhibited more rapid EGFR degradation than shC cells (Fig. 9C-F). These results suggest that α2,6 sialylation of EGFR protects against degradation following EGFR activation.

**FIGURE 9.**
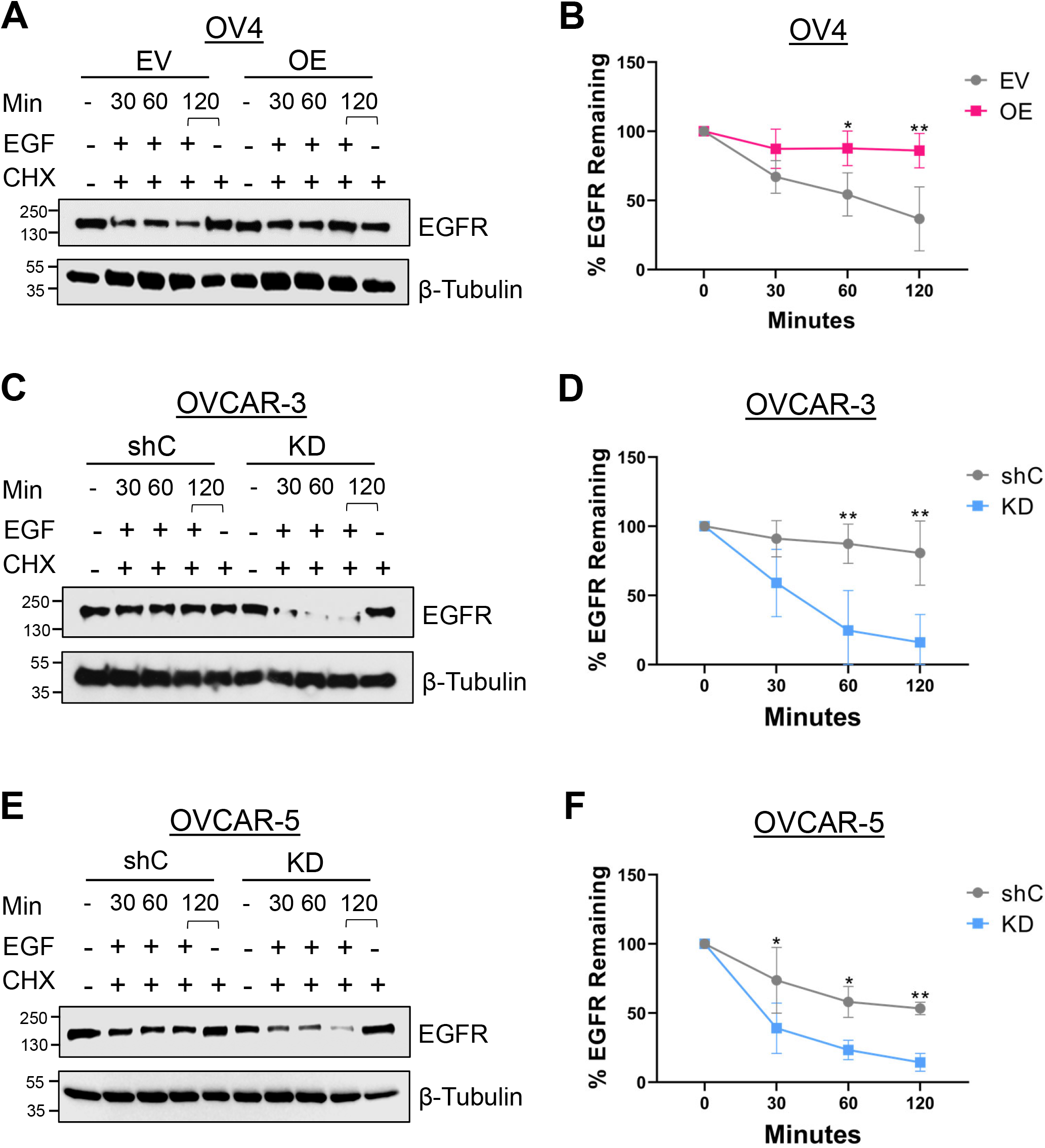
α2,6 sialylation of EGFR slows EGF-stimulated EGFR degradation. Cells were pretreated with 10 µg/mL of CHX for 2 hours to prevent nascent protein synthesis. Cells were then treated with EGF for 30, 60, or 120 minutes (in the continuous presence of CHX). As controls, cells were left untreated, or treated for 120 minutes with CHX alone. A-B). OV4 cells: representative immunoblots (A) and quantification (B) of the percent EGFR remaining. C-D). OVCAR-3 cells: representative immunoblots (C) and quantification (D) of the percent EGFR remaining. E-F). OVCAR-5 cells: representative immunoblots (E) and quantification (F) of the percent EGFR remaining. The percent EGFR remaining was calculated by densitometry, comparing levels of EGFR in the EGF-treated cells to the CHX controls. Graphs depict mean ± S.D. from three independent experiments. Statistics were calculated by using a two-way ANOVA followed by Šidák’s multiple comparison test (ns: p > 0.05, **: p < 0.01, ***: p < 0.001, ****: p < 0.0001).

### α2,6 sialylation of EGFR promotes higher-order EGFR clustering

To reinforce the biochemical assays described above, we evaluated EGFR activation and trafficking by microscopy. We utilized the OV4 cell model for these studies because OV4 OE and EV cells serve as an “on/off” system for ST6GAL1 expression (given that OV4 parental cells have no detectable endogenous ST6GAL1). Total Internal Reflection Fluorescence (TIRF) microscopy was used to assess higher-order clustering of EGFR, which has been proposed to promote EGFR activation and downstream signaling (35). TIRF selectively images within 100 nm of the cell membrane, making it an excellent method to study membrane protein distribution on the cell surface (36). TIRF was combined with Reflection Interference Contrast Microscopy (RICM), a method that detects the cell’s contact area with the surface of the coverslip, thus enabling measurements of the spread area of the adhered cell (representative RICM and TIRF images in Fig. 10A). RICM analyses showed that a 5-minute treatment with EGF stimulated cell spreading, and the cell contact area was larger in EGF-treated OE vs. EV cells (Fig. 10B). TIRF was then used to monitor EGFR clustering, and data were normalized to the cell contact area. Compared with EGF-treated EV cells, EGF-treated OE cells displayed a significant increase in the number (Fig. 10C) and size (Fig. 10D) of EGFR clusters, as well as an increase in the integrated surface EGFR intensity (Fig. 10E). No differences in EGFR clustering were noted in EV and OE cells in the absence of EGF simulation. These data support the hypothesis that α2,6 sialylation enhances EGFR homodimerization and higher-order clustering.

**FIGURE 10.**
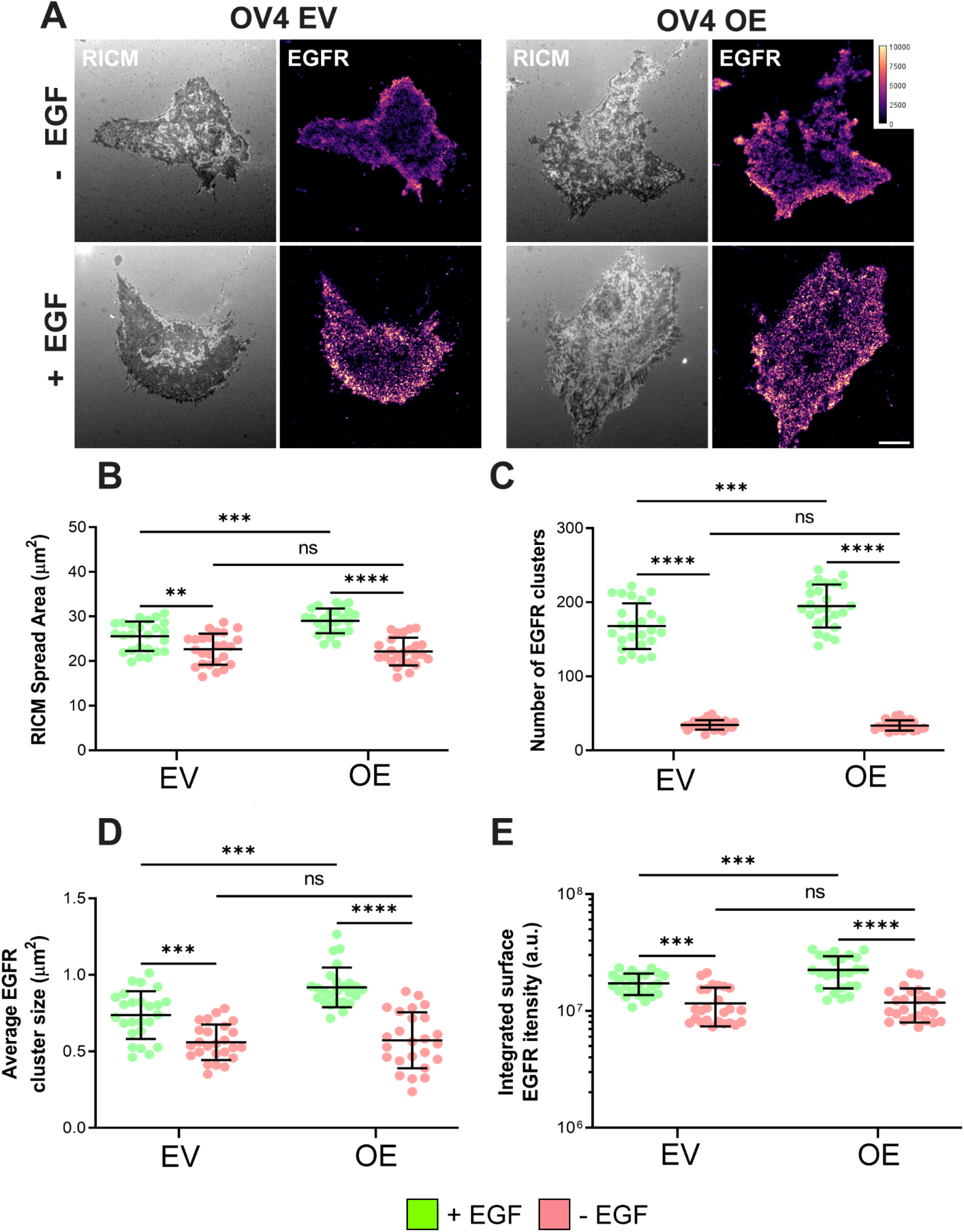
ST6GAL1-mediated EGFR sialylation promotes higher order EGFR clustering. A). OV4 EV and OE cells were treated with or without EGF for 5 minutes and stained for EGFR. Represent images are shown for cells visualized by RICM (grayscale) or TIRF. Images depict EGFR distributed on the plasma membrane, scale bar = 10 μm. B) RICM data showing the spread area of the cell. C-E) TIRF results (normalized to the area of the cell as indicated by RICM) with quantification of: the number of EGFR clusters per cell (C); the average EGFR cluster size (μm^2^) (D); and the integrated surface EGFR intensity (E). Graphs depict mean +/- S.D. from two independent experiments with 25 cells analyzed per group, per experiment. Data were analyzed by ANOVA with Tukey’s test, ns: p > 0.05, **: p < 0.01, ***: p < 0.001, ****: p < 0.0001.

### Upon EGF stimulation, α2,6 sialylated EGFR exhibits enhanced co-localization with recycling endosomes and decreased co-localization to lysosomes

To monitor EGFR trafficking throughout the cell, widefield z-stack images were acquired and deconvolved, allowing the generation of 3D reconstructions portraying EGFR localization within distinct subcellular compartments including endosomes and lysosomes. To assess recycling endosomes, cells were treated with EGF for 30 minutes and then co-stained for EGFR and Rab11, an established recycling endosomal marker (37). In agreement with the recycling assays shown in Fig. 8, we found that EGFR in OV4 OE cells had significantly greater colocalization with Rab11-positive endosomes following EGF treatment as compared with EV cells (representative images in Fig. 11A; quantification in Fig. 11B). To assess lysosomal co-localization, we treated cells with EGF for 60 minutes and co-stained cells for EGFR and the lysosomal marker, LAMP1 (38). In this case, OE cells had reduced co-localization of EGFR and LAMP1 compared with EV cells, suggesting decreased trafficking to the lysosome (representative images in Fig. 12A; quantification in Fig. 12B). Lysosomal-mediated degradation is the predominant mechanism by which EGFR is degraded (7); therefore these data align with results in Fig. 9 showing enhanced EGFR degradation in cells lacking ST6GAL1. Taken together, these data suggest that the α2,6 sialylation of EGFR acts as a switch to divert EGFR trafficking to recycling endosomes, thus promoting EGFR surface localization and downstream signaling.

**FIGURE 11.**
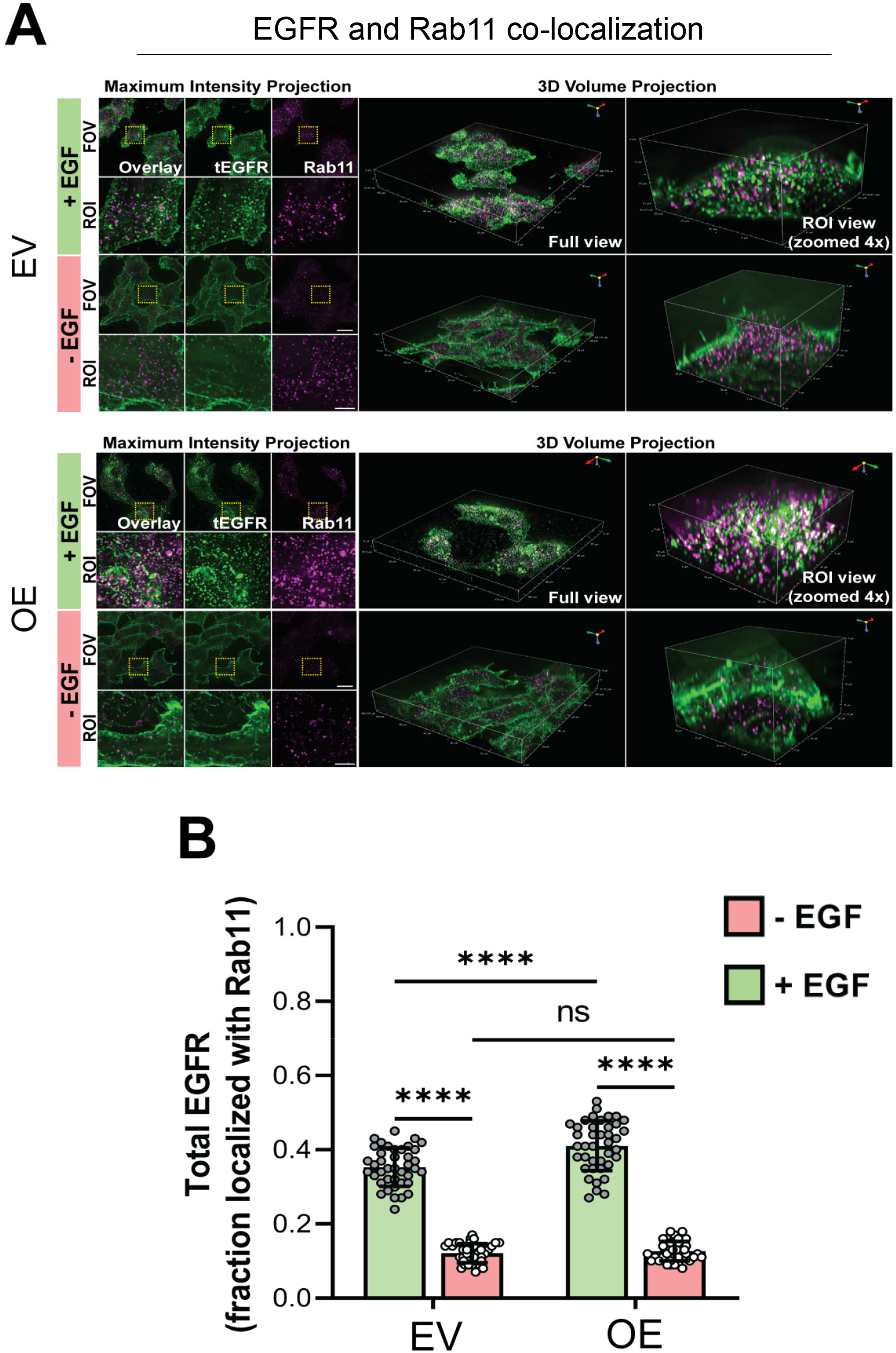
Sialylation of EGFR by ST6GAL1 promotes EGFR association with Rab11-positive recycling endosomes. A). Maximum intensity projection and 3D volume projection images for OV4 EV and OE cells treated with or without EGF for 30 minutes. Images were obtained using 3D widefield-deconvolution microscopy. The images depict the distribution of EGFR (green) and Rab11 (magenta) obtained following the processing of the acquired widefield 3D Z-stack images by the Richardson-Lucy algorithm for deconvolution. Scale bar for the field of view (FOV) = 20 μm, region of interest (ROI) = 5 μm. B). Quantification of the fraction of EGFR co-localized with Rab11-positive endosomes was executed using the JACoP plugin in Fiji. Graphs depict mean +/- S.D. from two independent experiments with 20 cells analyzed per group, per experiment. Data were analyzed by one way ANOVA with Tukey’s test (ns: p > 0.05, ****: p < 0.0001).

**FIGURE 12.**
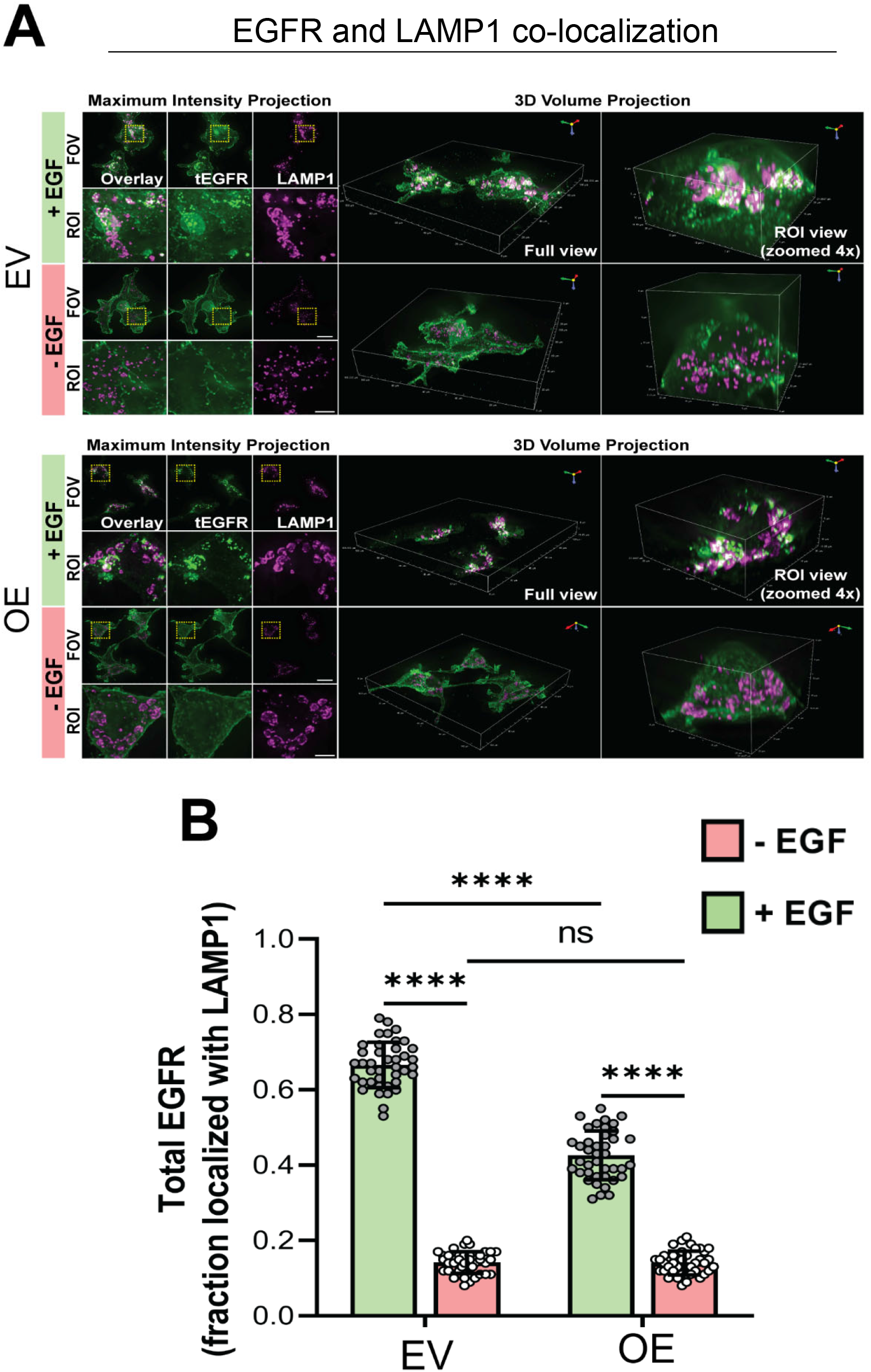
Sialylation of EGFR by ST6GAL1 reduces EGFR localization with LAMP1-positive lysosomes. A). Maximum intensity projection and 3D volume projection images for OV4 EV and OE cells treated with or without EGF for 60 minutes. Cells were visualized by 3D widefield-deconvolution microscopy. The images depict the distribution of EGFR (green) and LAMP1 (magenta) obtained following the processing of the acquired widefield 3D Z-stack images by the Richardson-Lucy algorithm for deconvolution. Scale bar for the field of view (FOV) = 20 μm, region of interest (ROI) = 5 μm. B). Quantification of the fraction of EGFR co-localized with LAMP1-positive lysosomes was executed using the JACoP plugin in Fiji. Graphs depict mean +/- S.D. from two independent experiments with 20 cells analyzed per group, per experiment. Data were analyzed by one way ANOVA with Tukey’s test (ns: p > 0.05, ****: p < 0.0001).

## Discussion

Alterations in glycosylation have long been associated with cancer (39, 40), however, compared with other areas of cancer research, cancer glycobiology remains greatly understudied. One of the predominant glycan changes in a cancer cell is an increase in α2,6-linked sialic acids on N-glycans, which occurs, in part, as a consequence of ST6GAL1 upregulation (18–21). ST6GAL1-mediated sialylation imparts pro-tumorigenic properties by modulating the structure and function of select cell surface receptors (20, 41). For instance, ST6GAL1-mediated sialylation of the TNFR1 and Fas death receptors prevents ligand-induced apoptosis by hindering receptor internalization (42–45), an event required for caspase activation. Additionally, α2,6 sialylation of CD45 and PECAM modulates receptor oligomerization (46, 47), whereas α2,6 sialylation of the β1 integrin promotes cell migration and invasion (48–50). Finally, we and others have identified EGFR as a target for ST6GAL1-mediated sialylation (22–27). However, the mechanisms by which α2,6 sialylation modulates EGFR activation and downstream signaling were previously unclear.

In the present study, we examined EGFR activation in cells with ST6GAL1 KD or OE, or in cells with high or low surface α2,6 sialylation as indicated by SNA staining. Across these various models, high ST6GAL1 expression and α2,6 sialylation consistently correlated with the activation of EGFR. Liu *et al.* described similar results in T-cell acute lymphoblastic leukemia cells, finding that ST6GAL1 KD diminished, and ST6GAL1 OE promoted, EGFR signaling (25). Other groups, however, have reported an inhibitory effect of sialylation on EGFR (26–29). Wong’s group showed that treatment of cancer cells with a sialidase enzyme caused an increase in EGFR activation, which was attributed to enhanced EGFR clustering (28, 29). However, the sialidase utilized in these studies cleaves all of the major sialic acid linkages (α2,3, α2,6, and α2,8). The broad ablation of sialoglycans from the cell surface is not biologically equivalent to selectively eliminating the α2,6 sialylation on N-glycans (18). In addition to Wong’s work, Park et al. (27) and Rodrigues et al. (26) reported a negative correlation between ST6GAL1 activity and EGFR activation. The reasons underlying the contradictory results regarding the effects of EGFR sialylation are not currently understood. One factor worth noting is that the SW48 cell line was used as a model in many prior studies that suggested an inhibitory effect of α2,6 sialylation (26, 27). SW48 cells harbor a G719S mutation in EGFR, which has been shown to promote ligand-independent activation of the receptor (30). Nonetheless, in our studies, the overexpression of ST6GAL1 in SW48 cells enhanced EGF-induced EGFR activation, consistent with our other cell models. While additional research will be needed to address the discrepant results regarding ST6GAL1’s effects on EGFR, we find that ST6GAL1 activity activates EGFR in the seven cell models studied herein, in addition to four other cell models described in our prior publications (22–24). Moreover, EGFR is markedly activated in the acinar cells of transgenic mice with forced expression of ST6GAL1 in the pancreas (51).

Our studies further show that the ST6GAL1-mediated sialylation of EGFR promotes formation of the active EGFR homodimer, as well as higher order clustering of EGFR. Other investigators have assessed the effects of global sialylation on EGFR dimer formation and clustering (28, 29, 52), however our results highlight a critical function for a specific sialic acid linkage, mediated by a unique sialyltransferase, in regulating EGFR dimerization and oligomerization. In addition, we demonstrate that α2,6 sialylation modulates the trafficking and fate of EGFR following EGF-induced receptor internalization. Results from recycling assays and 3D Z-stack imaging indicate that sialylation of EGFR by ST6GAL1 promotes its recycling and association with Rab11-positive recycling endosomes. Correspondingly, α2,6 sialylation of EGFR inhibits its degradation and association with LAMP1-positive lysosomes. Prior studies have reported that glycosylation modulates EGFR degradation (53–55), however, the effect of ST6GAL1-mediated sialylation on EGFR degradation was previously unexplored. Likewise, this is the first report demonstrating a role for α2,6 sialylation in EGFR trafficking, to our knowledge.

It is well known that the glycosylation of EGFR plays a pivotal part in regulating its structure. For example, the N-glycan on Asn-579 is critical for the formation of the auto-inhibitory tether. Ablation of this N-glycan weakens the tether, enabling the assembly of pre-formed dimers in the absence of ligand (17). Reis’ group reported that the Asn-579 N-glycan is, in fact, sialylated in cells with ST6GAL1 overexpression (Asn-579 is listed as Asn-603 in this reference due to the inclusion of the signal peptide in amino acid numbering) (26). It is tempting to speculate that the addition of the bulky, negatively-charged sialic acid to the Asn-579 N-glycan might interfere with formation of the auto-inhibitory tether, promoting EGFR activation. Like the Asn-579 site, an N-glycan on Asn-420 helps to maintain an inactive EGFR conformation. Deletion of the Asn-420 N-glycan promotes spontaneous oligomer formation and constitutive EGFR activation (56). In other studies, molecular dynamics simulations have indicated that N-glycans form noncovalent interactions with amino acids in the EGFR extracellular domain, which, in turn, stabilizes the EGF binding site (15). Finally, the N-glycosylation of EGFR contributes to the orientation of the EGFR ectodomain (16). In particular, the EGFR N-glycans adjacent to the plasma membrane help propel the ligand binding domains I and III away from the membrane, thereby facilitating EGF binding. These various investigations underscore the importance of N-glycans in regulating EGFR structure and activation, however the specific role of sialylation in these processes remains undetermined.

Beyond modulating EGFR signaling, it has been reported that the α2,6 sialylation of EGFR promotes resistance to various types of EGFR-targeted therapies such as the tyrosine kinase inhibitor (TKI) gefitinib, and the monoclonal antibody, cetuximab (23, 26, 27). Hence, it is essential to understand the mechanisms by which sialylation of EGFR regulates its structure and function. The current investigation shows that ST6GAL1-mediated sialylation of EGFR promotes receptor dimerization, clustering and recycling, thereby slowing EGFR degradation and promoting pro-survival signaling through AKT and NFκB. EGFR recycling, as well as signaling by AKT and NFκB, play well-known roles in fostering resistance to radiotherapy and also targeted therapies including antibodies and tyrosine kinase inhibitors (57–59). Our collective results provide novel insights into the functional consequences of EGFR sialylation in regulating its activation, signaling networks, and trafficking dynamics in malignant cells.

### Experimental procedures

#### Cell culture

MiaPaCa-2, OVCAR-3, and OVCAR-5 cells were obtained from ATCC. S2-013 cells were donated by Dr. Michael Hollingsworth at the University of Nebraska (Omaha, NE). OV4 cells were obtained from Dr. Timothy Eberlein at Harvard University (Cambridge, MA). S2-LM7AA cells were donated by Dr. Donald Buchsbaum at the University of Alabama at Birmingham (Birmingham, AL). Cells were grown in DMEM (MiaPaCa-2), RPMI-1640 (OVCAR-3, OVCAR-5, Suit-2, S2-013, S2-LM7AA), Leibovitz-L15 (SW48) or DMEM/F12 (OV4) supplemented with 1% antibiotic/antimycotic supplements (Gibco, 15240-062). OVCAR-3 cells were supplemented with 20% FBS and 0.01 mg/mL of bovine insulin (Sigma, I0516) and all other cells were supplemented with 10% FBS. All cell lines were grown in 5% CO_2_ except for the SW48 line, which was grown in 0% CO_2_. SW48 and OV4 cells were transduced with lentivirus encoding an empty vector (EV) (Sigma) or the human ST6GAL1 gene (OE) (Genecopoeia). OVCAR-3, OVCAR-5, S2-013, S2-LM7AA cells were transduced with lentivirus containing a shRNA control sequence targeting GFP (shC) (Sigma) or shRNA against ST6GAL1 (KD) (Sigma, TRCN00000035432, sequence: CCGGCGTGTGCTACTACTACCAGAACTCGAGTTCTGGTAGTAGTAGCACACGTTTTTG). MiaPaCa-2 cells were transduced with an empty vector lentivirus (EV) or the above mentioned sequence for shRNA against the ST6GAL1 gene (KD). Transductions were all performed using an MOI of 5 and stable polyclonal populations were selected using puromycin (5 µg/mL). Modulation of ST6GAL1 expression was confirmed by SNA staining and immunoblotting. For EGF treatments, cells were serum-deprived for 2 hours using media with 1% FBS. 100 ng/mL of EGF was then added in 1% FBS containing media for the indicated time intervals (R&D Systems, 236-EG-01M).

#### Immunoblotting

Cells were treated with or without EGF followed by lysis in radioimmunoprecipitation assay (RIPA) buffer (Pierce, 89901) supplemented with protease and phosphatase inhibitors (Pierce, 78440). Total protein concentration was confirmed by BCA assay (Pierce, 23225). Proteins were resolved by SDS-PAGE, and transferred to a polyvinylidene difluoride membrane (Millipore, IPVH00010). Membranes were blocked in 5% nonfat dry milk in TBS containing 0.1% Tween-20 (TBS-T). Membranes were then probed with antibodies for t-EGFR (1:1000, Cell Signaling Technologies, 4267), p-EGFR (1:1000, pTyr1068, Cell Signaling Technologies, 3777), t-AKT (1:1000, Cell Signaling Technologies, 4691), p-AKT (1:1000, pSer473, Cell Signaling Technologies, 4060), t-NF-κB p65 (1:1000, Cell Signaling Technologies, 8242), p-NF-κB p65 (1:500, pSer536, Cell Signaling Technologies, 3033), t-ERK1/2 (1:1000, Cell Signaling Technologies, 4695) and p-ERK1/2 (1:1000, p-Thr202/p-Tyr204, Cell Signaling Technologies, 4377). Blots were washed and incubated in HRP-conjugated anti-rabbit secondary antibodies (1:2500, Cell Signaling Technologies, 7074). Equal protein loading was confirmed using β-tubulin (1:2500, Abcam, ab21058 or 1:1000, Invitrogen, MA5-16308). Blots were developed with ECL Western Blotting Substrate (Pierce, 32106), Clarity Western ECL Substrate (Bio-Rad, 1705061), or SuperSignal West Femto (Pierce, 34096). Blots are representative of at least three independent experiments. Densitometry was measured in Fiji (ImageJ, National Institute of Health), and the phospho-proteins were normalized to their respective total protein to get a relative densitometry value which was then normalized to β-tubulin, the loading control. All statistics were calculated in GraphPad Prism (Version 9.5.1) using a two-way ANOVA followed by Šidák’s multiple comparison test. All results are shown as the mean +/- the standard deviation (S.D.).

#### Flow cytometry

Adherent cells were detached with accutase (BioLegend, 423201) and blocked on ice with 1% BSA in PBS. Cells were washed with 0.01% BSA in PBS and incubated with their corresponding antibodies for 30 minutes on ice. For total EGFR surface staining, cells were stained with 10 µg/mL of EGFR-Alexa Fluor 488 Clone AY13 (Biolegend, 352908). For SNA staining, cells were incubated with 20 µg/mL of SNA-FITC (Vector, FL-1301-2). For p-EGFR/SNA co-staining, cells were treated with EGF for 10 minutes as described under “Cell culture”. After treatment with EGF, cells were washed, fixed in 3.7% paraformaldehyde (PFA) (Electron Microscopy Services, 15710) permeabilized in 0.003% (*v/v*) Triton X-100, washed in PBS and stained with p-EGFR (p-Tyr1068) antibody (Cell Signaling Technologies, 3777) at a 1:1000 dilution. Cells were simultaneously stained with 20 µg/mL SNA. Cells were washed in PBS and anti-rabbit Alexa Fluor 488 (Invitrogen, A-11034) was added at 4 µg/mL. After staining, cells were washed and evaluated on the LSRII flow cytometer (BD Biosciences). Data were analyzed using FlowJo version 8 software (BD Biosciences) to obtain the MFI. For p-EGFR/SNA analysis, the 10% of cells with the highest levels of SNA staining were designated as “SNA high” and the 10% of cells with the lowest levels of SNA staining were denoted as “SNA low”. Levels of p-EGFR staining were then measured in these populations. Statistics were performed using GraphPad Prism. A two-way ANOVA was used followed by Šidák’s multiple comparison test. Results shown represent the MFIs +/- S.D. from three independent experiments.

#### SNA lectin precipitation

500 µg of cell lysate was incubated with 150 µg of SNA-agarose on a rotator at 4⁰C overnight (Vector Labs, AL-1303). Proteins containing α2,6 sialic acid were precipitated by centrifugation and washed 3 times with ice-cold PBS. Precipitates were then immunoblotted for EGFR as described above.

#### Ligand binding assay

Cells were detached using accutase and blocked in 1% BSA on ice, as previously described. Cells were then incubated with serial dilutions ranging from 0.39 nM to 200 nM of biotin-conjugated EGF (Invitrogen, E3477) in 0.01% BSA for 1 hour on ice (to minimize EGF/EGFR internalization). The cells were washed with PBS and incubated in 1 µg/mL of streptavidin conjugated to Alexa Fluor 488 (Invitrogen, S11223) in 0.01% BSA for 30 minutes. Cells were analyzed via flow cytometry. To obtain the fraction of maximum staining, the MFI at each concentration was divided by the MFI at the highest concentration of EGF. Values were plotted against the log of the concentration used. Graphs represent the mean +/- S.D. from three independent experiments. Data were graphed in GraphPad Prism.

#### BS^3^ cross-linking

The cross-linking protocol was adapted from Turk et al. 2015 (32). Cells were treated with EGF at 37⁰C for the indicated times and then immediately placed on ice. Cells were washed with ice-cold PBS and BS^3^ (Pierce, PG82083) was subsequently added to a final concentration of 3 mM. Cells were incubated with BS^3^ on ice for 20 minutes and the reaction was quenched with 250 mM glycine for 5 minutes. Cells were washed with PBS and lysed. Lysates were immunoblotted for EGFR as above. Densitometry was employed to evaluate levels of the EGFR dimer and monomer, and data were reported as the dimer to monomer ratio. Data shown are from three independent experiments.

#### Recycling assay

Cells were incubated in 10 µg/mL of CHX (Sigma Aldrich, C7698) for 2 hours at 37⁰C and then placed on ice for 5 minutes. Media containing 1% FBS, 10 µg/mL CHX and 100 ng/mL of EGF was subsequently added and cells were incubated for 15 minutes on ice to allow EGF to bind EGFR. Cells were then switched to 37⁰C for 15 minutes to enable internalization of the EGF/EGFR complexes. Following this incubation, an aliquot of cells was fixed in PFA and stained for EGFR to obtain a baseline measurement of the amount of EGFR remaining on the surface after the internalization step (designated as “time 0” for the recycling assay). For the remaining cells, EGF-containing media was replaced with EGF-free media containing 1% FBS and 10 µg/mL CHX, and cells were incubated at 37⁰C for 60 minutes to allow receptor recycling. Cells were subsequently detached with accutase and fixed in 3.7% PFA. EGFR staining was performed as above. Percent recycling was calculated by subtracting the MFI at time 0 from the MFI obtained at 60 minutes. This value was then divided by the MFI at 60 minutes and multiplied by 100 to obtain a percentage value. Statistics were then calculated in GraphPad Prism using a Student’s t-test. Results are shown as the mean +/- S.D.

#### Degradation assay

Cells were pre-incubated in media containing 1% FBS and 10 µg/mL CHX for 2 hours. EGF was then added to the cells as previously described and incubated for 30, 60 or 120 minutes (CHX was continuously present in the media during EGF treatment). Cells were lysed and immunoblotted for EGFR. As controls, cells were either left untreated, for treated with CHX alone for 120 minutes. Densitometric values were calculated using ImageJ and normalized to β-tubulin. The percent EGFR remaining was calculated by comparing normalized densitometric values to the CHX control. Statistics were performed using GraphPad Prism using a two-way ANOVA followed by Šidák’s multiple comparison test. Data are plotted as the mean +/- S.D.

#### RICM and TIRF microscopy

OV4 EV and OE cells were seeded overnight on glass coverslips (Thorlabs, CG15XH) coated with fibronectin (Sigma, F1141). Cells were serum-starved (1% FBS) for 2 hours and then treated with EGF for 5 minutes as described under “Cell culture”. Cells were fixed using 3.7% formaldehyde (Electron Microscopy Services, 15710) for 10 minutes at 37⁰C. Cells were then washed with PBS 5 times, permeabilized, and blocked with 0.25% Triton X-100 and 1% BSA for 30 minutes. Cells were stained for 2 hours at 37⁰C with primary antibody against EGFR (1:50, Invitrogen, MA5-13269). Cells were then washed 5 times in PBS and incubated for one hour with secondary anti-mouse Alexa Fluor 488 (1:250, Invitrogen, A32766) 37⁰C. After washing in PBS, cells were imaged in FluoroBrite DMEM (Gibco, A1896701). To evaluate surface EGFR distribution and clustering, TIRF and RICM were conducted as previously described (60). Briefly, OV4 cells were imaged on a Nikon Eclipse Ti2 microscope using the Nikon Elements software with an oil immersion Apo TIRF 60× NA 1.49 objective and an ORCA-Flash 4.0 V3 Digital CMOS camera (Hamamatsu). The sample was illuminated with a Sola epifluorescence light source (Lumencor) for RICM or with 488 nm laser for TIRF.

#### 3D *widefield microscopy*

OV4 EV and OE cells were seeded overnight on glass coverslips coated with fibronectin as above. For Rab11 imaging, cells were treated with EGF at 37⁰C for 30 minutes and for LAMP1 imaging, cells were treated with EGF for 60 minutes. Cells were fixed and blocked as described under RICM and TIRF microscopy. Cells were stained for t-EGFR (1:50, Invitrogen, MA5-13269), Rab11 (1:50, Cell Signaling, 5589), or LAMP1 (1:100, Cell Signaling, 9091) for 2 hours at 37⁰C. Cells were then washed and incubated for 1 hour at 37⁰C with anti-rabbit Alexa Fluor 647 (1:250, Invitrogen, A32733) or anti-mouse Alexa Fluor 488 (1:250, Invitrogen, A32766). Cells were washed in PBS and imaged in FluoroBrite DMEM (Gibco, A1896701). To obtain widefield Z-stacks, cells were imaged on a Nikon Eclipse Ti2 microscope using Nikon Elements software with a 100 nm step size in the Z dimension. Images were acquired with a 470/40 excitation filter and a 525/50 emission filter for Alexa Fluor 488 or a 620/60 excitation filter and a 700/75 emission filter for Alexa Fluor 647.

#### Image processing and analysis

Custom-written ImageJ macros were employed to subtract background fluorescence and measure morphological parameters, including the area of the cell footprint (RICM area), integrated intensity, and size and number of EGFR clusters. The RICM image was outlined manually to define the cell boundary and calculate the cell area. Integrated EGFR intensity was determined by subtracting the background measured from an off-cell region and then calculating the total fluorescence intensity within the cell boundary. The number and size analyses for EGFR clusters were estimated using the analyze particle function in Fiji following default thresholding of the background-subtracted image to generate a mask. The widefield Z-stack images were deconvolved using Nikon Elements deconvolution software (Richardson Lucy; parameters: 50 iterations, low noise level). Colocalization analysis was performed using the Fiji (ImageJ, National Institute of Health) plugin JACoP (Just Another Co-localization Plugin) to quantify Mander’s correlation coefficients (61). Statistical analysis by one-way ANOVA was performed using GraphPad Prism. All results are presented as mean +/- S.D.

## Data availability

All data described in this study are contained within the manuscript or supplementary figure (Figure S1).

## Supporting information

This article contains supporting information.

## Acknowledgements

The authors would like to thank Jihye Hwang for technical assistance and the UAB Comprehensive Flow Cytometry Core.

