## Supplementary Figure 1 for "Sialylation of EGFR by ST6GAL1 induces receptor activation and modulates trafficking dynamics"

**Running title:** ST6GAL1 activates EGFR and modulates receptor trafficking

**Material Included:**

Methods for Supporting Information

Figure S1. Monensin treatment attenuates EGFR recycling in cells with high ST6GAL1 expression.

**Methods for Supporting Information**

*Recycling assay with monensin treatment*

Recycling assays were performed in the presence or absence of 10 µM monensin, a recycling inhibitor. Cells were first preincubated in media containing 10 µg/mL of cyclohexamine (CHX) (+/- monensin) for 2 hours at 37ºC to block nascent EGFR synthesis. For the recycling assay, cells were incubated for 15 minutes on ice with 100 ng/mL of EGF in media containing CHX (+/- monensin) to allow EGF to bind EGFR. Cells were then switched to 37ºC for 15 minutes to enable internalization of the EGF/EGFR complexes. Following this incubation, the EGF-containing media was replaced with EGF-free media containing CHX (+/- monensin) and cells were incubated at 37ºC for 60 minutes to allow receptor recycling. Cells were stained for surface EGFR and analyzed by flow cytometry as described in the manuscript. Histograms depict the amount of surface EGFR at the end of the 60-minute recycling interval in the presence or absence of monensin.

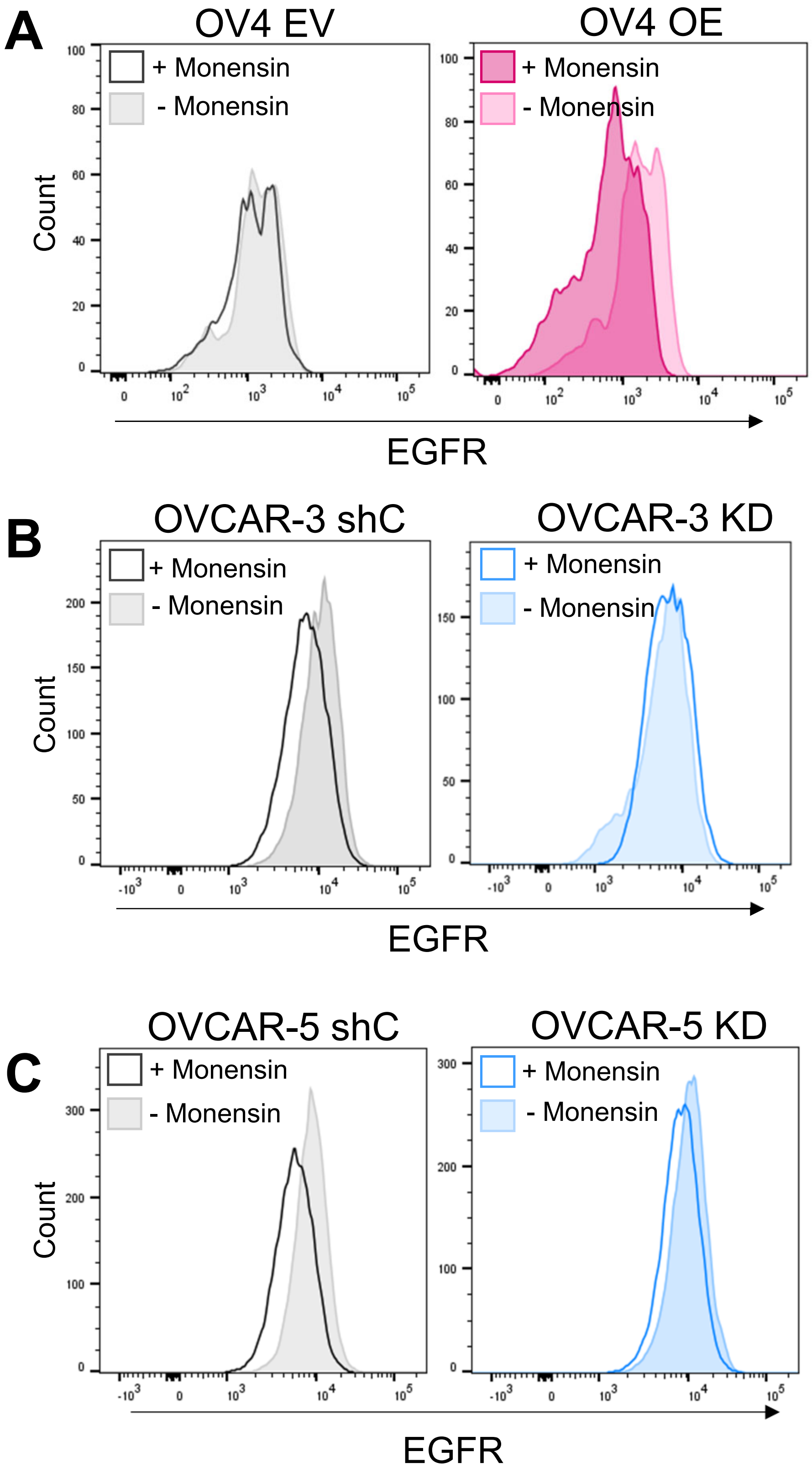


**Figure S1: Monensin treatment attenuates EGFR recycling in cells with high ST6GAL1 expression.** Recycling assays were performed in the presence or absence of 10 μM monensin, as described in the supplementary methods. After a 60 minute recycling interval, the amount of EGFR on the cell surface was measured by flow cytometry. A-C) Histograms depict Mean Fluorescent Intensity (MFI) for surface EGFR in OV4 cells (A), OVCAR-3 cells (B) and OVCAR-5 cells (C). Monensin treatment appeared to have a greater effect on cells with high expression of ST6GAL1, consistent with a positive role for sialylation in EGFR recycling.
